# Primary and secondary auditory cortex connectivity with brain regions involved in cognitive and emotional processing: in mouse and human

**DOI:** 10.1101/2025.06.26.661219

**Authors:** Büşra Köse-Özkan, Mazhar Özkan, Yasin Celal Güneş, Oktay Algın, Safiye Çavdar

## Abstract

Auditory cortex connectivity extends beyond the processing of acoustic stimuli, playing a crucial role in cognitive and emotional regulation through its interactions with higher-order brain regions. Although the neural mechanisms underlying acoustic information processing along the auditory pathway are well-documented, the connections supporting auditory-related cognitive and emotional processing, particularly in comparative studies between mice and human adults, are not yet fully clarified. In this study, we aim to investigate connections between the auditory cortex and brain regions involved in cognitive and emotional processing using retrograde fluoro-gold (FG) tracer in mice and 3-tesla high-resolution diffusion tensor tractography (DTI) in human adults. The FG injections into the primary (AI)/ secondary (AII) auditory cortices showed afferent connections with cortical (olfactory bulb, piriform, orbitofrontal, cingulate, motor, primary somatosensory, insular, visual, parietal, entorhinal and perirhinal cortices), subcortical (amygdala, hippocampus, globus pallidus, claustrum, bed nucleus of stria terminalis, diagonal band of the Broca and medial septal nucleus) and brainstem (raphe nuclei, pedunculopontine nucleus and locus coeruleus) structures. The DTI data obtained from human adults mostly corresponded with the experimental findings. Auditory cortical processing integrates auditory signals with other sensory, limbic and motor inputs. The connections collectively may suggest its role in cognitive and emotional functions. The auditory cortex is likely a critical hub within the neural circuitry underlying multisensory integration, decision-making, prediction, learning and memory functions. Understanding the connectivity of the auditory cortex can deepen our insight into its contribution to cognitive/emotional functions, offering new perspectives on the underlying mechanism linking hearing deficits with cognitive/emotional disorders.

## Introduction

Auditory information processing depends on neuronal physiology, the organization of neural networks, and the way these circuits transform signals throughout the auditory pathway. (Winer, 2010). The auditory cortex, corresponding to Brodmann areas 41, 42, and 22, is located in the temporal lobe of the human brain (Ong & Garey, 1990). Research in non-human primates and post-mortem human studies indicates that the auditory cortex is generally composed of core, belt, and parabelt areas, which are distinguished according to their micro-anatomical properties, functional features and connectivity (Hackett et al., 1998; Morosan et al., 2005). Typically, the core area, which consists of the primary auditory cortex (AI), is surrounded by and connected to other regions, often called secondary or belt areas (AII) (Kanwal & Ehret, 2010). Additionally, associative and multimodal fields in the parabelt (tertiary) area are involved in the integration of cognitive and acoustic representations of sound stimuli (Munoz-Lopez et al., 2010).

In the AI, one of the higher stages of the auditory hierarchy, tonotopic organization continues to play a central function, with sound representation mainly driven by acoustic properties. However, at this level, non-acoustic features in the AI increasingly modulate neuronal responses, influencing a range of aspects from the sequential arrangement of acoustic information to decision-making processes. Additionally, in secondary auditory areas outside the core region, neural activity may correspond more closely to perception rather than sensation (Nelken, 2020).

Auditory experiences such as exposing to noise or auditory deprivation have been shown to induce structural and functional alterations in the auditory pathway and also influence brain regions involved in cognitive and emotional processing (Kraus & Canlon, 2012; Zhang et al., 2021; Zhou & Merzenich, 2012). Epidemiological studies have provided evidence that environmental noise exposure has adverse effects on children’s cognitive functions and learning abilities (Lercher et al., 2003; Stansfeld et al., 2005). Hearing deficits such as hearing loss, tinnitus or impaired spectral and temporal processing are found to be linked to reduced performance on assessments of global cognition (Loughrey et al., 2018), executive functioning (Gates et al., 2010), attention (Shinn-Cunningham & Best, 2008), learning (Iliadou et al., 2009), memory (Köse et al., 2022; Rönnberg et al., 2011), speed of processing (Lee, 2018), and anxiety/depression (Hackenberg et al., 2023). Growing evidence suggests that these cognitive and emotional impairments are underpinned by disruptions in the connectivity between auditory cortical regions and higher-order cognitive networks (An et al., 2023; De Ridder et al., 2022; Xu et al., 2019). Disruptions to these pathways, whether through peripheral hearing damage or prolonged sensory deprivation, may thus play a pivotal role in the emergence of cognitive decline and altered emotional processing associated with hearing deficits.

The neural mechanisms underlying the processing of acoustic information along the pathway to the auditory cortex are well documented, however, the connections supporting auditory cognition and emotion processing, as well as the distinctions between the AI and AII, have yet to be fully understood, particularly given the connectivity differences between the human and mouse brain. The present study aimed to define the connections of the AI and AII with brain regions involved in both cognitive and emotional processing using the neuroanatomical tract-tracing technique in mice and three-tesla DTI in healthy human subjects.

## Materials and Methods

### 2.1. Ethics Statement

Mice were housed in an AAALAC International-accredited animal facility and the Institutional Animal Care and Use Committee of Koç University have approved all procedures.

### 2.2. Animals

Data from sixteen successful injections with minimal contamination of adjacent structures out of twenty-one male C57BL/6J mice were included in the present study. Animals weighing 26-30 g were housed under a 12 h/12 h light/dark cycle in a room with constant temperature (20 ± 3°C) and allowed access to water and food *ad libitum*.

### 2.3. Fluoro-Gold procedures

To investigate the afferent connectivity of the AI and AII, retrograde tracer injections were performed in mice using Fluoro-Gold (FG). FG was dissolved in the distilled water to yield a 5% concentration. Mice were anesthetized with ketamine (90 mg/kg, intraperitoneally) and xylazine (10 mg/kg, intraperitoneally) based on their body weights. Animals were secured in a stereotaxic frame (Stoelting Quintessential Injector, Wood Dale, IL, USA), and a midline scalp incision was made to expose the skull between bregma and lambda. A small craniotomy was performed over the skull at a position appropriate for the unilateral injection of FG into the right AI (n=6) or AII (n=10). A Hamilton Syringe (Hamilton 1701, 32 ga, 2 in, point style 3) containing 0.1 µL FG solution (5%, FluoroChrome Inc. Englewood, CO, USA) was lowered into AI (AP: −3.07 ± 0.2 mm, ML: 4 ± 0.2 mm, DV: 0.5 ± 0.2 mm) and AII (AP: −2.03 ± 0.2 mm, ML: 4 ± 0.2 mm, DV: 0.7 ± 0.2 mm) unilaterally (Fig. 1A, B, C and D), according to the mouse brain atlas (Paxinos & Franklin, 2012). After a 7-day survival period, mice were deeply anesthetized with a higher dose of ketamine (300 mg/kg, intraperitoneally) and xylazine (30 mg/kg, intraperitoneally) and perfused transcardially with phosphate-buffered saline solution, followed by 4% paraformaldehyde in 0.1 M phosphate buffer. Brains were extracted, post-fixed overnight at 4°C in the same fixative, and sectioned coronally at 40 μm thickness using a cryostat (Leica, Heidelberger, Germany). Every third section was mounted onto a glass slide, dehydrated, coverslipped with anti-fade fluorescence mounting medium (ab104136, Abcam), and examined under a fluorescence microscope (Zeiss, Jena, Germany). The density of ipsilateral and contralateral FG labeled cells and axons were qualitatively categorized as follows: *sparse: for cell numbers 1–5, **moderate: for cell numbers 6–20 ***dense: for cell numbers above 20. Notably, we do not report on the connections between AI and AII with regions within the central auditory pathway, as these were beyond the scope of this study and have been extensively covered previously in the literature (Budinger et al., 2000a, 2000b; Malmierca & Ryugo, 2010).

**Fig. 1.**
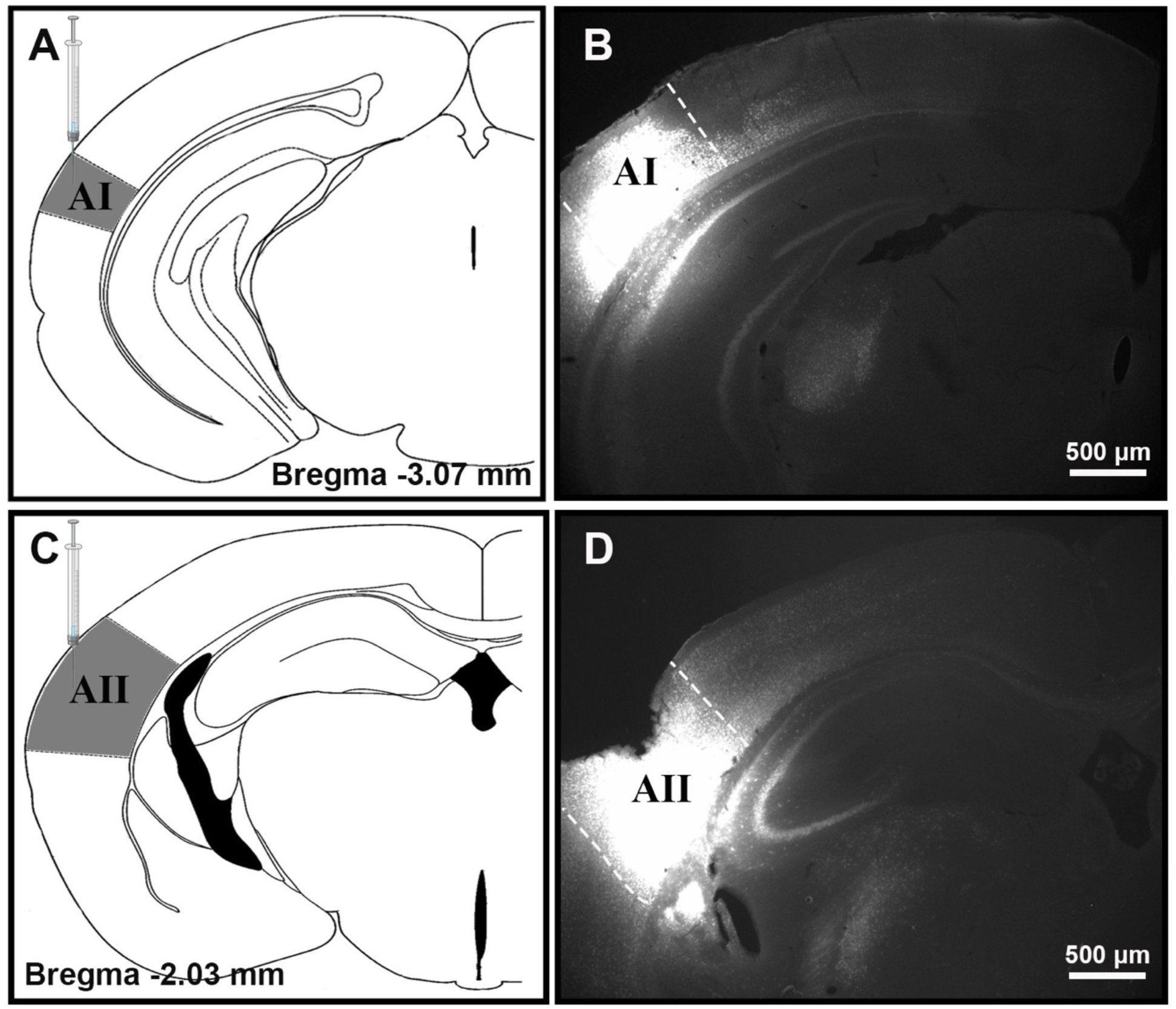
Injection sites in primary (AI) and secondary (AII) auditory cortices. Schematic illustrations showing the injection sites of Fluoro-Gold (FG) tracer into the AI and AII, respectively (A,C). Representative photomicrographs demonstrating FG injection sites in the AI and AII, respectively (B,D).

### 2.4. Diffusion Tensor Imaging (DTI)

The Human Connectome Project (HCP), conducted by the MGH-USC partnership, generated the DTI data. This dataset comprises 3T magnetic resonance (MR) scans of healthy individuals, including 19 males and 16 females, with an age range of 20–59 years and a mean age of 31 years. The scans were performed using the 3T Connectome scanner (MAGNETOM Skyra, Siemens Healthcare, Erlangen, Germany) at the Athinoula A. Martinos Center for Biomedical Imaging, located in Boston, Massachusetts, USA. Data acquisition utilized a 64-channel birdcage head coil, and all participant images were collected in NIFTI format. The MR examinations employed high-resolution sequences, including T2-weighted 3D-SPACE, T1-weighted 3D-MPRAGE, and diffusion-weighted imaging (DWI) acquired with four distinct b-values. The following parameters were utilized in the DWI sequence: field of view (FOV): 140 x 140 mm; matrix: 210 x 210; repetition time (TR)/echo time (TE): 8800/57 ms; acquisition time: 89 ms; b-values (s/mm^2^): 1000 (64 directions), 3000 (64 directions), 5000 (128 directions), and 10000 (256 directions); number of slices: 96; slice thickness: 1.5 mm; echo spacing: 0.63 ms; parallel imaging acceleration factor (iPAT): 3; and multiband factor.

The data were pre-processed using FreeSurfer V5.3.0 and FSL V5.0 programs. The Eddy tool was employed to correct distortions caused by eddy currents. The Harvard Ascending Arousal Network Atlas (AAN) was utilized to illustrate the connections between the auditory cortex and brain regions involved in cognitive and emotional processing. The AAL2-AAL3 and HCP 1065 atlases were used to depict the AI and AII, as well as regions involved in the cognitive and emotional processing of the human brain (http://brain.labsolver.org/diffusion-mri-templates/hcp-842-hcp-1021, https://surfer.nmr.mgh.harvard.edu/fswiki/CorticalParcellation, http://www.gin.cnrs.fr).

The Harvard AAN atlas illustrated the relationship between fibers originating from the auditory cortex and various structures. Seed sites were implanted to reveal the pathways connecting the auditory cortex to cognitive regions. The DSI Studio application was employed for data processing and tract visualization. By superimposing the previously segmented areas in the atlas with the images from the HCP dataset, proper synchronization was achieved.

## 3. Results

The connections of the AI and AII with cortical, subcortical, and brainstem regions involved in cognitive and emotional processing in mice were revealed using FG and in humans using DTI. The experimental studies in mice and DTI studies in humans demonstrated that AI and AII show both ipsilateral and bilateral connections with various brain regions. The contralateral connections passed within the corpus callosum to reach the contralateral brain areas. The density of FG-labeled neurons in cortical, subcortical, and brainstem structures involved in cognitive and emotional processing was documented (Table 1).

**Table 1.** The density of ipsilateral and contralateral Fluoro-Gold labeled neurons in cortical, subcortical, and brainstem structures was estimated as follows: * sparse: for the cell number 1-10, ** moderate: for cell number between 11-30, *** dense: for the cell number between 31-above in a specific area.

|  | Primary Auditory Cortex |  | Secondary Auditory Cortex |  |
| --- | --- | --- | --- | --- |
|  | ipsilateral | contralateral | ipsilateral | contralateral |
| Cortical |  |  |  |  |
| • Olfactory bulb (G1) | - | - | * | - |
| • Piriform cortex (Pir) | - | - | * | - |
| • Orbitofrontal cortex (OFC) | ** | * | *** | * |
| • Cingulate cortex (A24, A30, A32) | ** | * | ** | * |
| • Motor cortex (M1, M2) | * | - | *** | * |
| • Somatosensory cortex (S1) | * | - | ** | * |
| • Insular cortex (ICx) | * | - | ** | - |
| • Visual cortex (V1, V2) | ** | * | *** | ** |
| • Parietal cortex (PCx) | - | - | ** | * |
| • Entorhinal cortex (Ent) | - | - | * | * |
| • Perirhinal cortex (PRh) | - | - | * | * |
| Subcortical |  |  |  |  |
| • Amygdala (BLA) | *** | - | * | - |
| • Hippocampus | *** | ** | - | - |
| • Globus pallidus (GP) | ** | - | * | - |
| • Claustrum (Cl) | * | - | * | - |
| • Bed nucleus of stria terminalis (BNST) | * | - | - | - |
| • Diagonal band of the Broca (VDB, HDB) | * |  | - |  |
| • Medial septal nucleus (MS) | ** |  | * |  |
| Brainstem |  |  |  |  |
| • Raphe nuclei (DR, MnR) | * | - | - | - |
| • Pedunculopontine nucleus (PPN) | * | - | - | - |
| • Locus coeruleus (LC) | * | - | - | - |

### 3.1. The connections of the AI with brain regions involved in cognitive and emotional processing

In mice, AI demonstrated connections with various cortical, subcortical, and brainstem regions. The distinct bilateral cortical connections of the AI were with the primary and secondary visual (V1 and V2), orbitofrontal (OFC), and cingulate (A24, A30, A32) cortices. Furthermore, AI showed ipsilateral projections to the insular (ICx), primary somatosensory (S1) and primary and secondary motor (M1, M2) cortices. The DTI results in humans confirmed the cortical connections of the AI with ipsilateral visual, insular, lateral orbitofrontal cortices and isthmus of the cingulate cortex. However, DTI results did not show connections with the primary somatosensory cortex (S1) and motor cortex (M1, M2), which were observed in mice. The majority of the cortical connections in mice were bilateral; however, in humans, they were ipsilateral, and no bilateral cortical connections were observed.

The subcortical connections of the AI involved in cognitive and emotional processing were with basolateral amygdala (BLA) (Fig. 2B), hippocampus (CA3) (Fig. 3A, B), globus pallidus (GP), claustrum (Cl), bed nucleus of stria terminalis (BNST), diagonal band of the Broca (VDB, HDB) and medial septal nucleus (MS) (Fig. 4B). All subcortical connections of the AI were ipsilateral in mice, except those with the hippocampus (Table 1). Labeled intrahippocampal axons were observed in the ventral hippocampal commissure (vhc, Fig. 3C). The results of AI-subcortical connectivity using DTI confirmed the presence of bilateral connections with the amygdala (Fig. 2C), and the hippocampus (Fig. 3D), as well as the ipsilateral connections with the septal nucleus (Fig. 4C), and the bed nucleus of the stria terminalis. However, DTI results did not show connections with globus pallidus (GP), claustrum (Cl), or diagonal band of Broca, which were observed in mice.

**Fig. 2.**
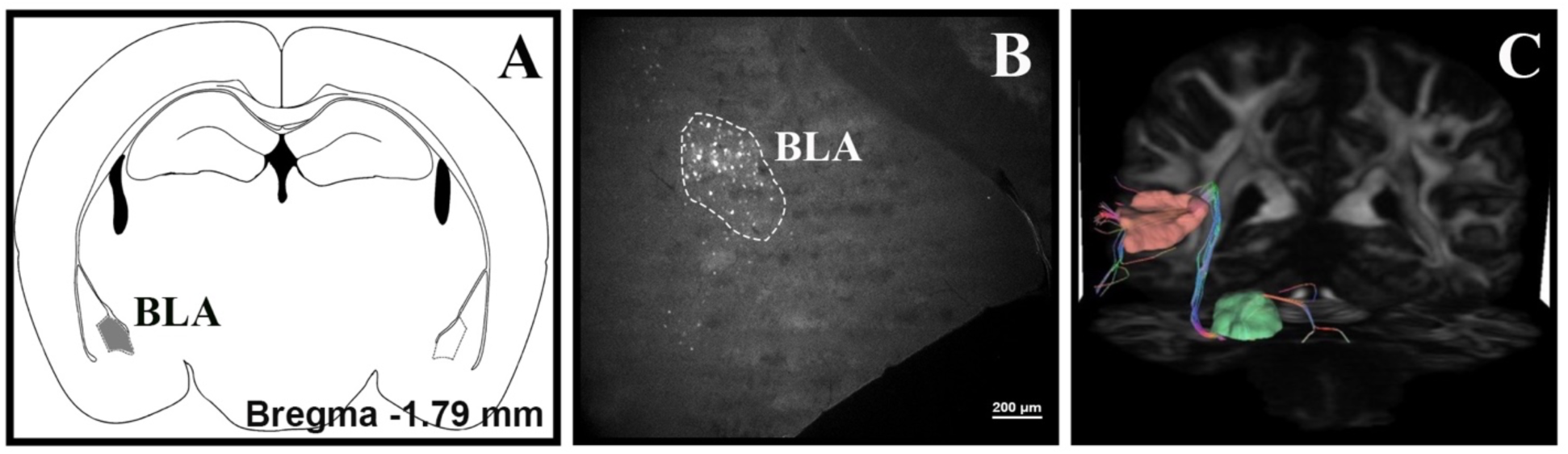
Ipsilateral FG labeling in basolateral amygdaloid nucleus (BLA) in mice and DTI-based connectivity between primary auditory cortex (AI) and the bilateral amygdala in humans. Schematic illustration showing the location of the BLA in the mouse brain atlas, provided for anatomical orientation of the FG-labeled region (A). FG-labeled neurons were demonstrated in the ipsilateral BLA (B). DTI analysis revealed bilateral connectivity between the AI and the amygdala; the representative image displays the tract on the ipsilateral side (C).

**Fig. 3.**
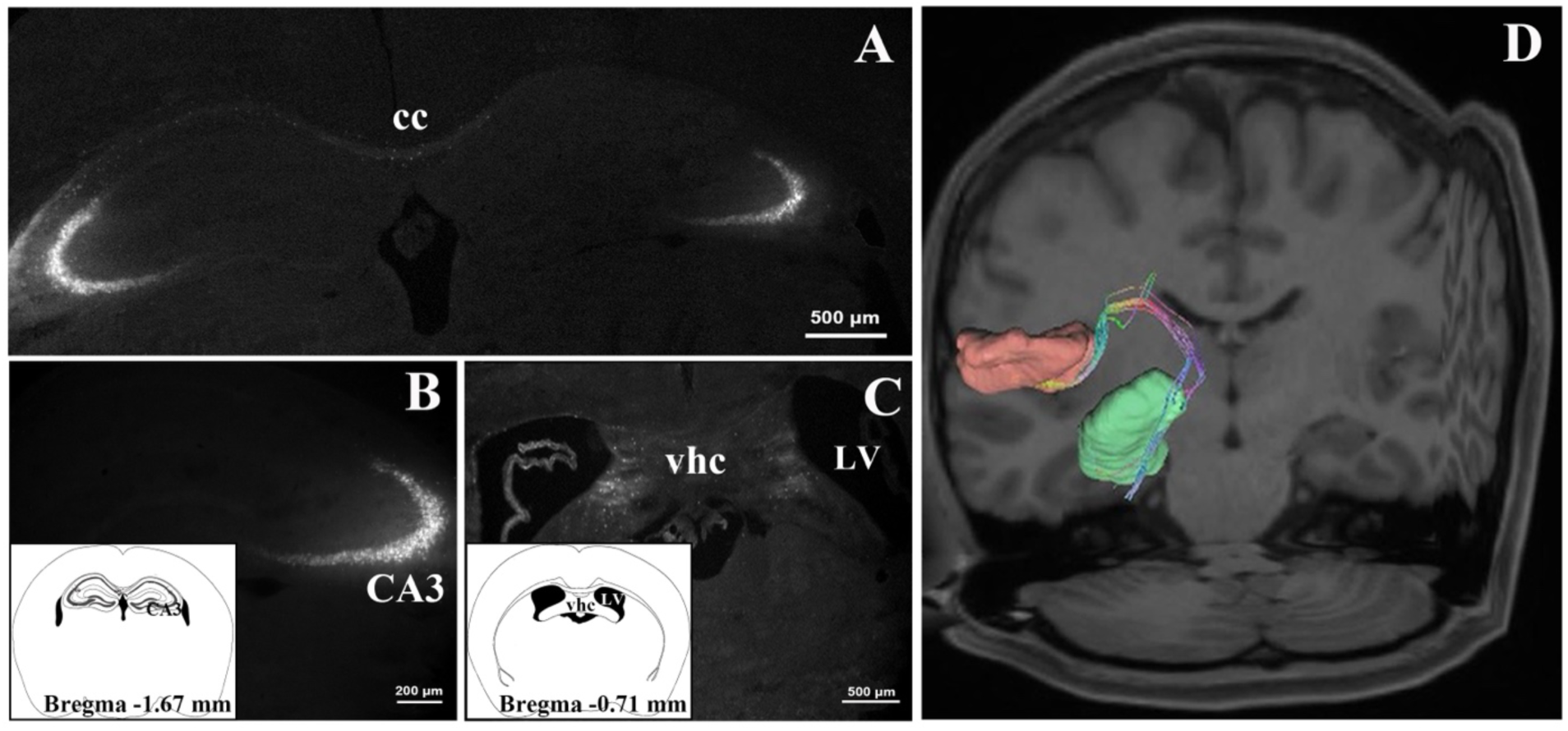
Bilateral FG labeling in CA3 and ventral hippocampal commissure in mice and DTI-based connectivity between primary auditory cortex (AI) and the bilateral hippocampus in humans. Bilateral FG-labeled neurons were detected in the field CA3 of the hippocampus (A, B) and ventral hippocampal commissure (vhc) (C). Schematic illustrations were added to the lower left corner for orientation. The DTI revealed connections between the AI and the bilateral hippocampus; the representative image displays the tract on the ipsilateral side (D). (cc, corpus callosum; LV, lateral ventricle)

**Fig. 4.**
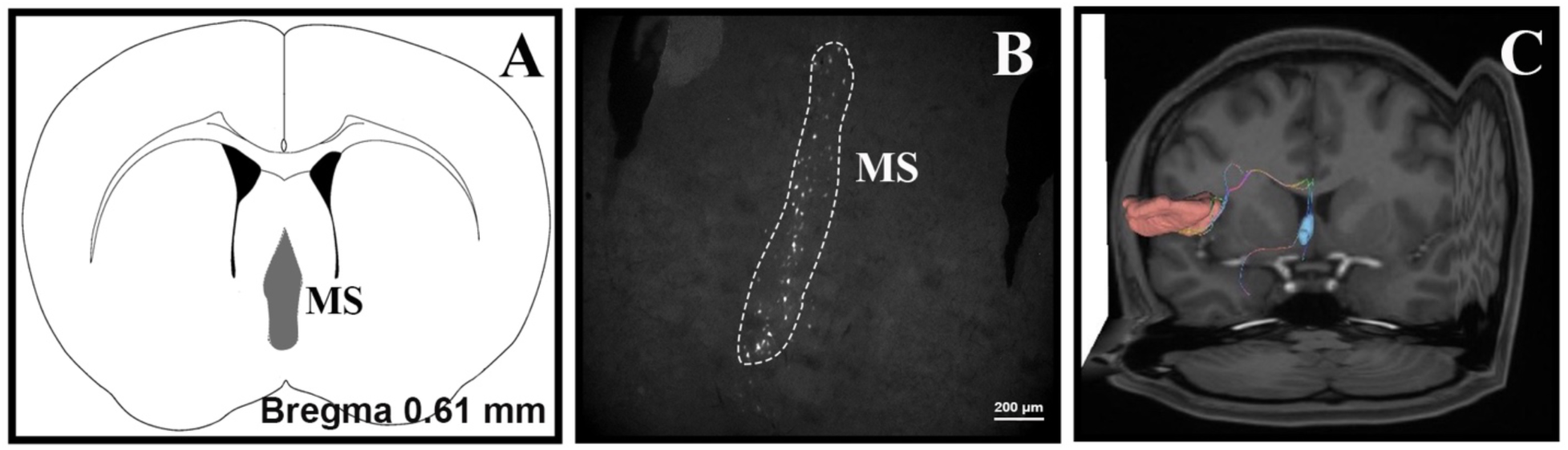
FG labeling in medial septal nucleus (MS) in mice and DTI-based connectivity between primary auditory cortex (AI) and the ipsilateral septal nucleus in humans. Schematic illustration showing the location of the MS in the mouse brain atlas, provided for anatomical orientation of the FG-labeled region (A). FG-labeled neurons were demonstrated in the MS (B). The DTI showed ipsilateral connections between the AI and the septal nucleus (C).

Following FG injections into the AI, labeled neurons were observed in brainstem structures. The AI received afferent fibers from the ipsilateral raphe nuclei (DR, MnR), pedunculopontine nucleus (PPN) and locus coeruleus (LC) (Table 1). The DTI results in humans confirmed the connectivity of the AI with locus coeruleus; no connections were observed with raphe nuclei (DR, MnR) and pedunculopontine nucleus (PPN). Additionally, DTI data revealed the presence of AI-rostral pons and AI-dentate nucleus connections in humans, which were not present in mice.

### 3.2. The connections of the AII with brain regions involved in cognitive and emotional processing

Injections of the retrograde tracer FG into the AII revealed afferent connections with cortical regions involved in cognitive and emotional processing. The distinct bilateral cortical connections of the AII were with the orbitofrontal (OFC) (Fig. 5B), cingulate (A24, A30, A32), primary and secondary motor (M1, M2) (Fig. 6B), primary somatosensory (S1) (Fig. 7B), primary and secondary visual (V1, V2) (Fig. 8B), parietal (PCx), entorhinal (Ent) and perirhinal (PRh) cortices. The bilateral connections with the cortical regions were dense on the ipsilateral side and sparse on the contralateral side. Additionally, ipsilateral connections were observed with the olfactory bulb (G1), piriform cortex (Pir) and insular cortex (ICx) (Table 1). The DTI results in humans confirmed the cortical connections of the AII with ipsilateral lateral orbitofrontal (Fig. 5C), entorhinal, perirhinal, parietal, insular, primary motor (Fig. 6C), primary somatosensory (Fig. 7C) and primary visual (Fig. 8C) cortices. However, DTI in human subjects did not show connections with the secondary motor (M2) and visual cortices (V2). The DTI results in humans exhibited the connectivity of the AII with bilateral caudal anterior (Fig. 9A) and isthmus of the cingulate cortex (Fig. 9B), while ipsilateral connections were observed with the posterior cingulate cortex (Fig. 9C). Whereas the majority of AII connections were bilateral in mice, all cortical connections in humans were ipsilateral, except those with the cingulate cortex. However, DTI results did not show connections with the olfactory bulb and piriform cortex, which were observed in mice.

**Fig. 5.**
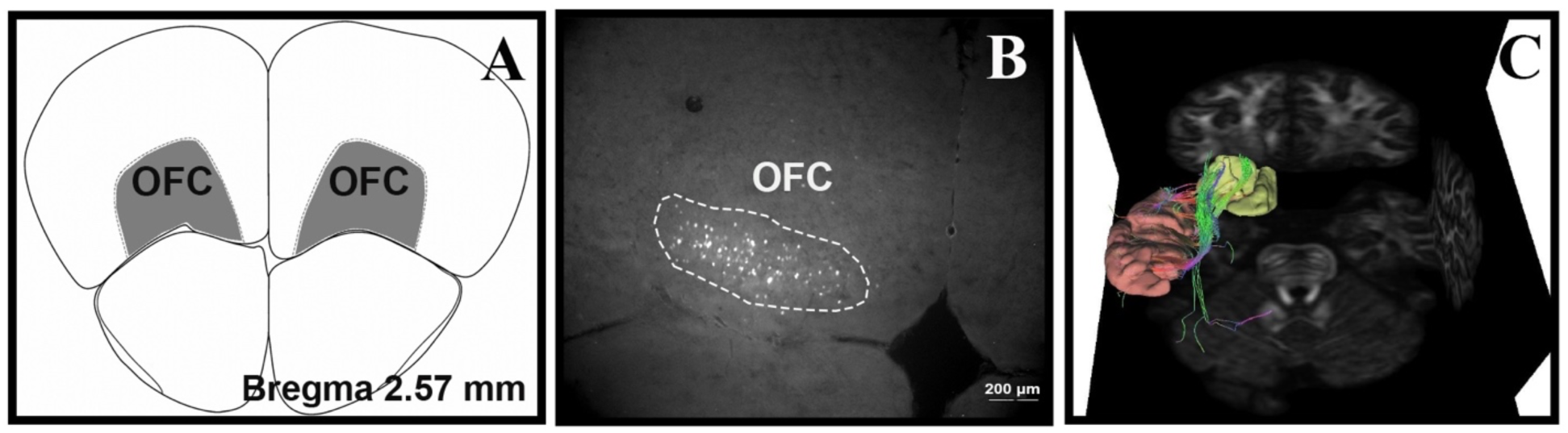
Bilateral secondary auditory cortex (AII) - orbitofrontal cortex (OFC) connectivity in mice and ipsilateral connections in humans. Schematic illustration showing the location of the OFC in the mouse brain atlas, provided for anatomical orientation of the FG-labeled region (A). FG-labeled neurons were observed in the bilateral OFC; the representative image displays labeling on the ipsilateral side (B). The DTI showed ipsilateral connections between the AII and the lateral OFC (C).

**Fig. 6.**
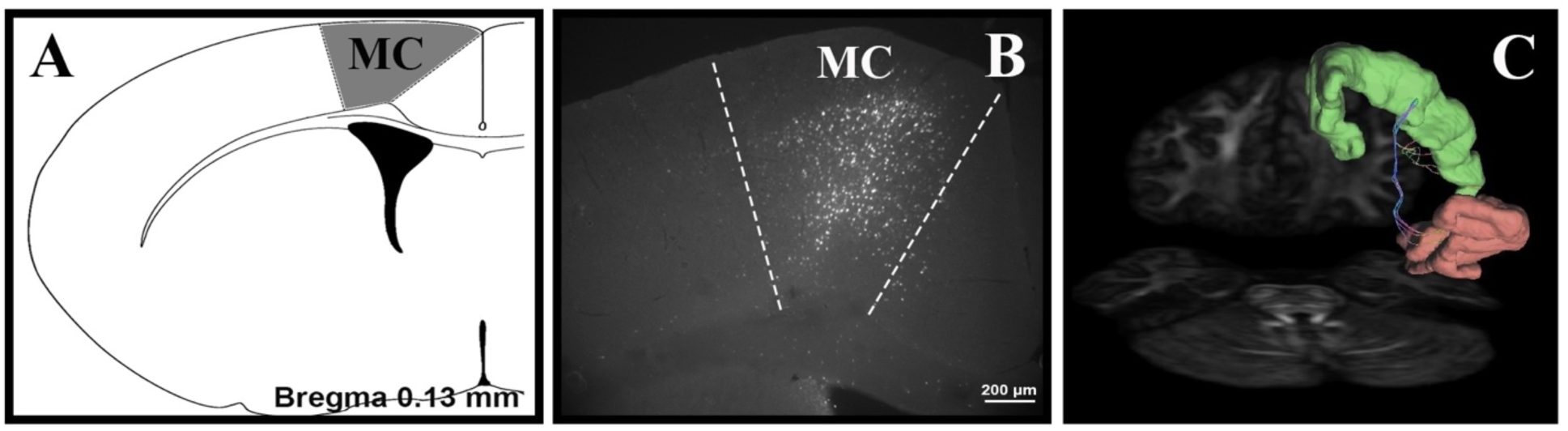
FG labeling in the primary and secondary motor cortex (MC) in mice and DTI-based connectivity between secondary auditory cortex (AII) and ipsilateral primary motor cortex in humans. Schematic illustration showing the location of the MC in the mouse brain atlas, provided for anatomical orientation of the FG-labeled region (A). FG-labeled neurons were detected in the MC (B). DTI data revealed the presence of AII-primary motor cortex (C) connections in humans.

**Fig. 7.**
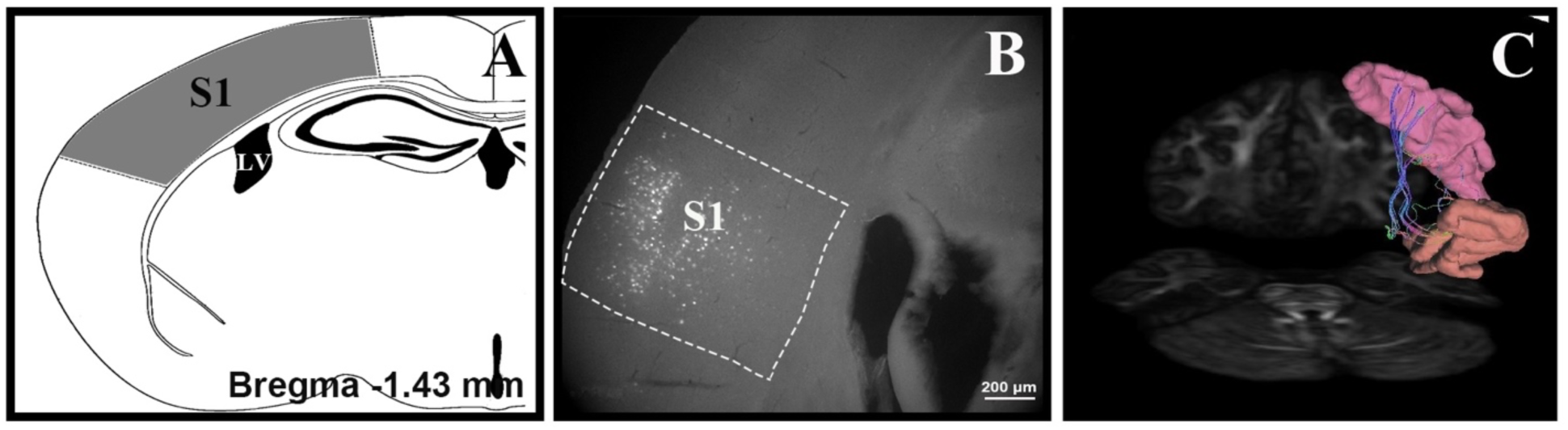
Bilateral FG labeling in the primary somatosensory cortex (S1) in mice and DTI-based connectivity between secondary auditory cortex (AII) and ipsilateral primary somatosensory cortex in humans. Schematic illustration showing the location of the S1 in the mouse brain atlas, provided for anatomical orientation of the FG-labeled region (A). FG-labeled neurons were revealed in the bilateral S1; the representative image displays labeling on the ipsilateral side (B). The DTI showed ipsilateral connections between the AII and the S1 (C).

**Fig. 8.**
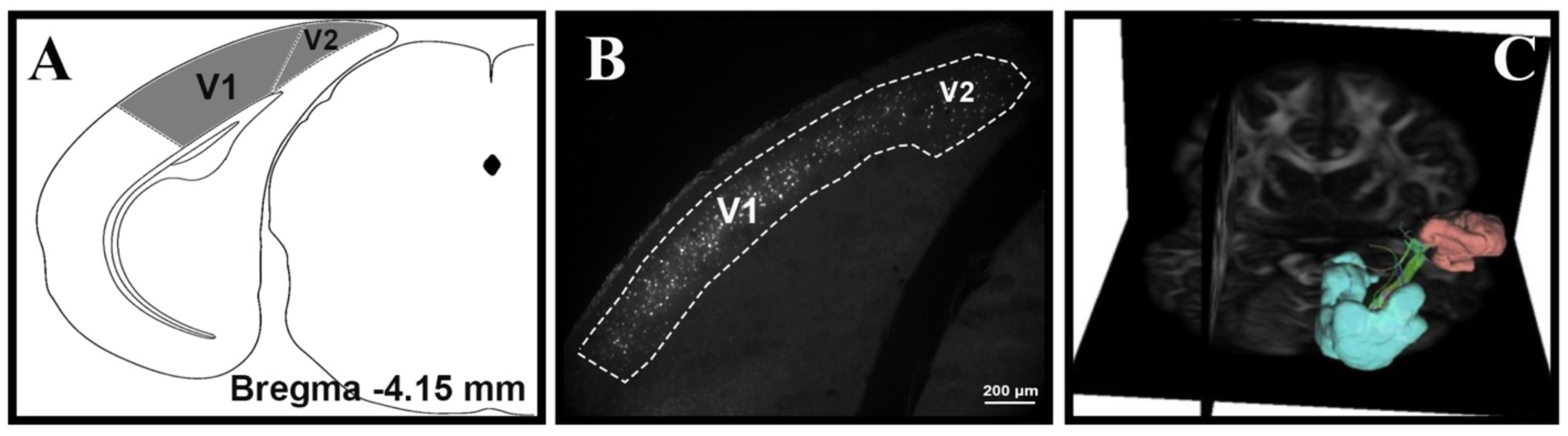
Bilateral FG labeling in the primary (V1) and secondary (V2) visual cortex in mice and DTI-based connectivity between secondary auditory cortex (AII) and ipsilateral primary visual cortex in humans. Schematic illustration showing the location of the V1 and V2 in the mouse brain atlas, provided for anatomical orientation of the FG-labeled region (A). FG-labeled neurons were detected in the V1 and V2 (B). The DTI showed ipsilateral connections between the secondary auditory cortex (AII) and the primary visual cortex (C).

**Fig. 9.**
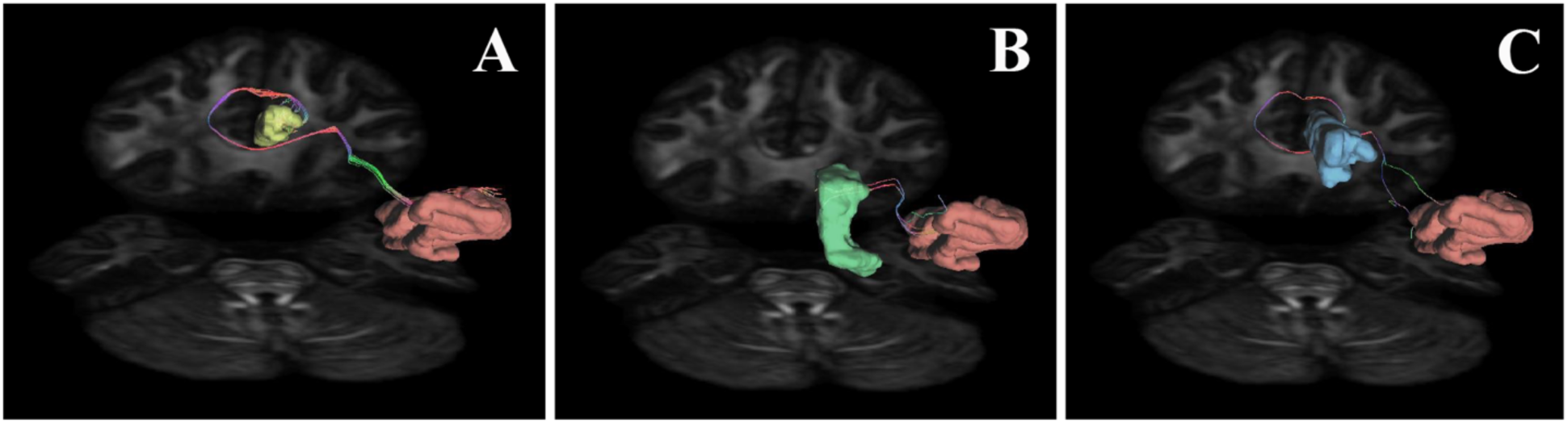
DTI revealed ipsilateral and bilateral connections between AII and subregions of the cingulate cortex in humans. DTI analysis revealed bilateral connectivity between the secondary auditory cortex (AII) and the caudal anterior (A) and isthmus of the cingulate cortex (B); the representative images display the tracts on the ipsilateral side. Also, ipsilateral connections were observed with the posterior cingulate cortex (C).

The subcortical connections of the AII in the mice included the ipsilateral amygdala, globus pallidus (GP), claustrum (Cl) and medial septal nucleus (MS). Consistent with the FG results, DTI in human subjects demonstrated ipsilateral connections between the AII and the amygdala and claustrum. However, DTI results did not show connections with the globus pallidus, which were observed in mice. In addition, DTI showed contralateral connections with the septal nucleus, while ipsilateral connections were not detected. AII showed no connections with brainstem structures in the mice; however, DTI data in humans revealed connections with the pedunculopontine nucleus (PPN).

## 4. Discussion

The connections between the auditory cortex and the central auditory pathway have been well documented (Pickles, 2015). However, the connections of the auditory cortex with cognition-related brain regions in Mongolian gerbils, monkeys, mice, and humans have been documented in limited detail (Budinger et al., 2008; Jang & Choi, 2022; Kraus & Canlon, 2012; Upadhyay et al., 2007; Yukie, 2002). In the present study, comprehensive data were gathered on the connectivity of the AI and AII with cognition-related brain regions. Furthermore, these connections were systematically compared between mice and humans.

### 4.1. Major findings

The present tract-tracing study shows ipsilateral or bilateral connections of the AI with cortical (orbitofrontal, cingulate, motor, somatosensory, insular and visual cortices), subcortical (amygdala, hippocampus, globus pallidus, claustrum, bed nucleus of stria terminalis, diagonal band of the Broca and medial septal nucleus) and brainstem (raphe nuclei, pedunculopontine nucleus, locus coeruleus) structures. Moreover, FG injections into the AII revealed inputs from cortical (olfactory bulb, piriform, orbitofrontal, cingulate, motor, somatosensory, insular, visual, parietal, entorhinal and perirhinal cortices) and subcortical areas (amygdala, globus pallidus, claustrum and medial septal nucleus). No connections were observed between AII and brainstem structures in mice. Subcortical structures associated with cognitive and emotional processing exhibited stronger connectivity with AI, while cortical areas were more strongly connected to AII.

The majority of the experimental findings in mice were confirmed by DTI in humans. However, the cortical connections of the AI, including those with the primary somatosensory cortex (S1) and primary and secondary motor (M1, M2), and the olfactory-associated cortical connections of the AII, including the olfactory bulb and piriform cortex, were not observed in humans. AII showed bilateral connections with the primary and secondary motor (M1, M2) and visual cortex (V1, V2) in mice; however, DTI in human subjects demonstrated ipsilateral connections with the primary motor (M1) and primary visual cortices (V1). Additionally, connections of the AI and AII with the globus pallidus, and the AI with the claustrum and the diagonal band of Broca, were not detected in humans, although they were present in mice. Brainstem connections of the AI with the raphe nuclei and pedunculopontine nucleus (PPN), as shown in mice, were also not detected in humans. Conversely, AI connections with the rostral pons and dentate nucleus and AII connections with the PPN were detected in humans but not in mice.

### 4.2. Cortical connections

Recent research on the auditory cortex has highlighted its critical role in the complex integration of multimodal and multisensory information, as well as in the evaluation of physical parameters specific to auditory stimuli (Chang et al., 2025; Solyga & Keller, 2025). Studies have reported that neuronal responses in the auditory cortex can be elicited or modulated by stimuli from other modalities, such as somatosensation (Brosch et al., 2005; Stehberg et al., 2014; Wallace et al., 2004) and vision (Bizley et al., 2007; Brosch et al., 2005). The connections between the auditory cortex and other sensory regions in both mice and humans may underlie mechanisms of multisensory integration and attentional modulation. Furthermore, we also identified projections from the olfactory bulb and piriform cortex to the AII. This finding corresponds with recent electrophysiological studies (Vogler et al., 2024; Wesson & Wilson, 2010). Wesson and Wilson identified auditory-responsive neurons in the olfactory tubercle, suggesting convergence of auditory and olfactory signals in subcortical regions (Wesson & Wilson, 2010). However, AI/AII– olfactory connections have not been identified in humans, which is likely attributable to species-specific differences in cortical organization and the relative reliance on olfactory sensory modalities between mice and humans.

Auditory processing also involves interactions with the non-sensory network of the forebrain (Hackett, 2011). The findings of the present study demonstrate that AI receives inputs from the orbitofrontal, insular, cingulate and motor cortices, whereas the AII additionally receives projections from the parietal, entorhinal, and perirhinal cortices. Previous optogenetic and electrophysiological studies have demonstrated that the OFC responds to both unisensory (auditory or visual) and multisensory (audiovisual) stimuli and projects to the auditory cortex, forming excitatory synapses on both pyramidal neurons and inhibitory interneurons across all layers of the AI (Sharma & Bandyopadhyay, 2020; Winkowski et al., 2018). Similarly, the insular cortex shows multisensory responsiveness; its caudal portion, in particular, may specialize in processing auditory information and contribute to the perception of communication sounds, such as speech (Remedios et al., 2009; Rodgers et al., 2008).

The cingulate cortex plays a central role in cognitive control, emotional regulation, and attentional processing (Bush et al., 2000; Vogt, 2009). In our study, while most cortical connections of the AII in humans were ipsilateral, DTI revealed bilateral connectivity specifically with the caudal anterior and isthmus regions of the cingulate cortex. The bilateral cingulate-AII connectivity in humans may reflect the critical role of the cingulate cortex in coordinating emotionally and cognitively relevant auditory processing across hemispheres, particularly during tasks that involve auditory attention, decision-making, or motivational salience. In accordance with the present findings, a recent behavioral study identified a top-down modulatory circuit from the cingulate cortex to the auditory cortex, enhancing auditory perceptual performance under challenging listening conditions (Anbuhl et al., 2025).

Motor cortex activity modulates auditory cortex processing by inducing global inhibition, which alters the neural state to enhance signal-to-noise ratio, sharpen frequency tuning, and adjust auditory responsiveness depending on behavioral context (Reznik & Mukamel, 2019). The absence of detectable connections between the auditory cortex and the M2 in humans, which were present in mice, may reflect differences in functional cortical organization across species. In contrast to mice, in humans, motor-induced shifts in auditory processing and auditory-guided motor behavior may be distributed across more complex networks, including parietal and premotor cortices (Hickok & Poeppel, 2007; Zatorre et al., 2007).

The extensive projections to the auditory cortex may contribute to higher-order processes, including perceptual decision-making, associative memory formation, and multisensory integration. In the present study, we have shown the connections of AII with entorhinal and perirhinal in both mice and humans. In this respect, entorhinal and perirhinal inputs may facilitate cortical plasticity and memory encoding (Li et al., 2014). Also, we have demonstrated the connections of AII with the parietal cortex in both mice and humans. Animal studies indicate that the parietal cortex and auditory cortex are interconnected to facilitate decision-making (Nelson & Mooney, 2016; Yao et al., 2020; Zhong et al., 2019). Silencing these parietal-to-auditory cortex projections impairs the animals’ ability to categorize new auditory stimuli, suggesting a direct influence of the parietal cortex on auditory processing (Zhong et al., 2019). Yao et al. further demonstrated that the parietal cortex plays a causal role in integrating auditory features for perceptual judgments. Inactivating the parietal cortex in gerbils led to increased minimum integration times during auditory discrimination tasks, indicating its involvement in auditory decision-making (Yao et al., 2020). Additionally, the motor cortex and basal forebrain provide converging movement-related inputs to the auditory cortex. While basal forebrain cholinergic input rapidly modulates auditory tuning, motor cortical input signals impending movements and motor planning (Nelson & Mooney, 2016). This convergence of bottom-up and top-down signals allows the auditory cortex to dynamically adjust sensory representations in real time, thereby supporting stable perception while enabling rapid adaptation to change behavioral and emotional contexts.

The bilateral AII-cortical connectivity observed in mice contrasts with predominantly ipsilateral connections in humans, which may be related to species-specific differences or the different techniques used to reveal the connections. In rodents, sensory cortices, including auditory, visual, and somatosensory areas, exhibit a high degree of interhemispheric connectivity, primarily mediated by the corpus callosum and anterior commissure (Olavarria & van Sluyters, 1985; Wise & Jones, 1976). These commissural pathways facilitate the bilateral integration of sensory information, which is essential for spatial localization, threat detection, and coordinating behavior in small mammals (Kaas, 2008). In contrast, the human cerebral cortex exhibits a more pronounced functional and structural lateralization, which is considered a fundamental feature of human brain organization and a “leading principle of human evolution” (Labache et al., 2023). The human cerebral cortex is more functionally and structurally lateralized, particularly in higher-order association areas such as those involved in language, attention, and emotion (Karolis et al., 2019; Labache et al., 2023; Toga & Thompson, 2003).

### 4.2. Subcortical connections

Acoustic inputs are known to induce the function of a wide range of subcortical structures. The hippocampus plays a well-established role in memory formation, perception, and cognition (Billig et al., 2022). A recent study demonstrated that during the mental replay of auditory sequences, information flows from the hippocampus to the auditory cortex, suggesting an active role in reconstructing auditory experiences from memory (Dimakopoulos et al., 2022). Moreover, the hippocampus can modulate auditory cortical responses, indicating its influence on the perception and interpretation of sound (Leong et al., 2023). This is consistent with the previous studies reporting that neurons in the limbic system, including the hippocampus, amygdala, cingulate cortex, striatum and septal nucleus, responded to noise burst stimuli and clicks (Bordi & LeDoux, 1992; Chen et al., 2014; Vinogradova, 1975; Xiao et al., 2018). In the current study, both mice and humans showed bilateral hippocampus-AI connectivity. This bilateral organization may reflect the hippocampus’s role in complex cognitive operations such as memory retrieval and memory-guided attention (Dimakopoulos et al., 2022; Teki et al., 2012).

The projections from the amygdala to the auditory cortex may play a key role in modulating auditory processing during emotionally significant experiences such as fear learning and memory (Tsukano et al., 2019). In the present study, mice showed ipsilateral connections, whereas bilateral connections between the amygdala and AI were observed in humans, suggesting a potentially more integrated or distributed mechanism of emotional modulation of auditory processing may be present in humans. Furthermore, the bilateral organization may enhance the brain’s capacity to coordinate affective auditory responses across hemispheres, particularly in complex social contexts and musical emotion discrimination (Frühholz et al., 2016; Manno III et al., 2019; Plichta et al., 2011).

Notably, in the present study, we identified a previously unreported connection between the BNST and AI in both mice and the human brain. Projections from both the amygdala and BNST may contribute to the emotional modulation of auditory processing, prioritizing or suppressing sensory inputs depending on affective states such as fear, anxiety, or depression (Lv et al., 2024; Wang et al., 2021). Additionally, the claustrum has been implicated in auditory scene analysis by reflecting sensory changes (Remedios et al., 2014). This may suggest that the claustrum contributes to processing complex auditory information, potentially through its connections with the auditory cortex (Chia et al., 2020).

A study has shown that cholinergic neurons in the basal forebrain show a high degree of sensory modality specificity in their projections to the auditory cortex (Kim et al., 2016). In the present study, we observed that AI receives inputs from cholinergic structures such as the diagonal band of Broca in mice and the septal nuclei in both mice and humans. These inputs from basal forebrain structures to the AI may be involved in auditory processing and associative learning related to punishment (Chaves-Coira et al., 2018; Robert et al., 2021) and may also be critical for dynamically modulating cortical activity in response to auditory input. The DTI results in the present study did not show connections with the globus pallidus or diagonal band of Broca. In rodents, these structures may be more directly integrated into auditory-limbic-motor loops, whereas in humans, auditory influence on these structures may occur via indirect or polysynaptic pathways.

### 4.3. Brainstem and cerebellar connections

Studies have shown that serotonergic projections from the ipsilateral raphe nuclei are known to influence cortical excitability (Puig & Gulledge, 2011; Ren et al., 2019) and may play a modulatory role in auditory processing, particularly in relation to emotional and arousal states. In mice, AI– raphe connectivity may support rapid modulation of auditory processing under stress or affective states. However, our DTI results did not show connections with the raphe nuclei. In humans, serotonergic modulation of the auditory cortex may be via indirect projections.

Optogenetic studies have shown that activation of noradrenergic neurons in the locus coeruleus induces widespread desynchronization across multiple sensory cortices (Carter et al., 2010; Kim et al., 2016). Specifically, noradrenergic projections from the locus coeruleus to the AI may be critical for regulating sleep-wake transitions and maintaining arousal-dependent auditory responsiveness (Kim et al., 2016).

Both anatomical tract-tracing and DTI data revealed that the PPN sends afferent projections to the auditory cortex, targeting the AI in mice and the AII in healthy human subjects. These findings suggest a potential modulatory role of the PPN in auditory cortical processing, likely mediated via its cholinergic output (Özkan et al., 2022). Furthermore, DTI-based evidence of AI connectivity with the rostral pons and dentate nucleus of the cerebellum indicates that the auditory cortex is embedded within a broader sensorimotor-integrative framework in humans. These brainstem and cerebellar pathways may contribute to functions such as speech production, audiomotor coordination, and the temporal organization of auditory-driven behaviors, highlighting the role of the auditory cortex in higher-order sensorimotor integration (Benarroch, 2024; Küper et al., 2011; Perales et al., 2006).

## Limitations

Although only successful and well-confined injections were included in the tract-tracing experiments, the potential for contamination of adjacent brain regions cannot be entirely excluded. The DTI remains the only non-invasive and safe method for assessing white matter connectivity in living human subjects; however, it presents several limitations. Because DTI infers structural connectivity based on the diffusion of water molecules rather than the active transmission of neural signals, it does not permit the determination of the directionality (afferent/efferent) of fiber tracts. Despite these limitations, the integration of DTI and neuroanatomical tracing provides complementary insights, and future studies can build on these findings using advanced imaging techniques to improve spatial and directional precision.

## Conclusion

The AI and AII receive inputs from the various brain regions involved in cognitive and emotional processing. Our results highlight both shared and species-specific features of auditory network organization. While mice exhibited bilateral cortical and subcortical connectivity, humans showed a greater degree of functional lateralization. The interspecies differences may reflect the increased specialization and hemispheric differentiation observed in the human brain, particularly for complex functions such as language, emotion regulation, and memory-guided attention.

Overall, our findings underscore the involvement of the auditory cortex in multisensory integration, emotional modulation, memory, and attentional control, extending far beyond classical auditory pathways. Understanding the connectivity of the AI and AII in both mice and the human brain will deepen our insight into their roles in cognitive and emotional functions, offering new perspectives on the mechanisms linking hearing deficits to cognitive and emotional disorders, and may also be critical for advancing targeted neurorehabilitation strategies.

## Acknowledgments

The authors would like to thank the Koç University Research Center for Translational Medicine (KUTTAM) for the use of the services and facilities. Also, DTI data were obtained from the Human Connectome Project.

## References

An, Y. Y., Lee, E.-S., Lee, S. A., Choi, J. H., Park, J. M., Lee, T.-K., Kim, H., & Lee, J. D. (2023). Association of hearing loss with anatomical and functional connectivity in patients with mild cognitive impairment. JAMA otolaryngology–head & neck surgery, 149(7), 571–578.

Anbuhl, K. L., Diez Castro, M., Lee, N. A., Lee, V. S., & Sanes, D. H. (2025). The cingulate cortex facilitates auditory perception under challenging listening conditions. Proceedings of the National Academy of Sciences, 122(14), e2412453122.

Benarroch, E. (2024). What Is the Role of the Dentate Nucleus in Normal and Abnormal Cerebellar Function? Neurology, 103(3), e209636.

Billig, A. J., Lad, M., Sedley, W., & Griffiths, T. D. (2022). The hearing hippocampus. Progress in neurobiology, 218, 102326.

Bizley, J. K., Nodal, F. R., Bajo, V. M., Nelken, I., & King, A. J. (2007). Physiological and anatomical evidence for multisensory interactions in auditory cortex. Cerebral cortex, 17(9), 2172–2189.

Bordi, F., & LeDoux, J. (1992). Sensory tuning beyond the sensory system: an initial analysis of auditory response properties of neurons in the lateral amygdaloid nucleus and overlying areas of the striatum. Journal of Neuroscience, 12(7), 2493–2503.

Brosch, M., Selezneva, E., & Scheich, H. (2005). Nonauditory events of a behavioral procedure activate auditory cortex of highly trained monkeys. Journal of Neuroscience, 25(29), 6797–6806.

Budinger, E., Heil, P., & Scheich, H. (2000a). Functional organization of auditory cortex in the Mongolian gerbil (Meriones unguiculatus). III. Anatomical subdivisions and corticocortical connections. European Journal of Neuroscience, 12(7), 2425–2451.

Budinger, E., Heil, P., & Scheich, H. (2000b). Functional organization of auditory cortex in the Mongolian gerbil (Meriones unguiculatus). IV. Connections with anatomically characterized subcortical structures. European Journal of Neuroscience, 12(7), 2452–2474.

Budinger, E., Laszcz, A., Lison, H., Scheich, H., & Ohl, F. W. (2008). Non-sensory cortical and subcortical connections of the primary auditory cortex in Mongolian gerbils: bottom-up and top-down processing of neuronal information via field AI. Brain research, 1220, 2–32.

Bush, G., Luu, P., & Posner, M. I. (2000). Cognitive and emotional influences in anterior cingulate cortex. Trends in cognitive sciences, 4(6), 215–222.

Carter, M. E., Yizhar, O., Chikahisa, S., Nguyen, H., Adamantidis, A., Nishino, S., Deisseroth, K., & De Lecea, L. (2010). Tuning arousal with optogenetic modulation of locus coeruleus neurons. Nature neuroscience, 13(12), 1526–1533.

Chang, S., Zheng, B., Keniston, L., Xu, J., & Yu, L. (2025). Auditory cortex learns to discriminate audiovisual cues through selective multisensory enhancement. Elife, 13, RP102926.

Chaves-Coira, I., Rodrigo-Angulo, M. L., & Nuñez, A. (2018). Bilateral pathways from the basal forebrain to sensory cortices may contribute to synchronous sensory processing. Frontiers in neuroanatomy, 12, 5.

Chen, G.-D., Radziwon, K. E., Kashanian, N., Manohar, S., & Salvi, R. (2014). Salicylate-Induced Auditory Perceptual Disorders and Plastic Changes in Nonclassical Auditory Centers in Rats. Neural plasticity, 2014(1), 658741.

Chia, Z., Augustine, G. J., & Silberberg, G. (2020). Synaptic connectivity between the cortex and claustrum is organized into functional modules. Current Biology, 30(14), 2777–2790. e2774.

De Ridder, D., Vanneste, S., Song, J.-J., & Adhia, D. (2022). Tinnitus and the triple network model: a perspective. Clinical and experimental otorhinolaryngology, 15(3), 205–212.

Dimakopoulos, V., Mégevand, P., Stieglitz, L. H., Imbach, L., & Sarnthein, J. (2022). Information flows from hippocampus to auditory cortex during replay of verbal working memory items. Elife, 11, e78677.

Frühholz, S., Trost, W., & Kotz, S. A. (2016). The sound of emotions—Towards a unifying neural network perspective of affective sound processing. Neuroscience & Biobehavioral Reviews, 68, 96–110.

Gates, G. A., Gibbons, L. E., McCurry, S. M., Crane, P. K., Feeney, M. P., & Larson, E. B. (2010). Executive dysfunction and presbycusis in older persons with and without memory loss and dementia. Cognitive and Behavioral Neurology, 23(4), 218–223.

Hackenberg, B., Döge, J., O’Brien, K., Bohnert, A., Lackner, K. J., Beutel, M. E., Michal, M., Münzel, T., Wild, P. S., & Pfeiffer, N. (2023). Tinnitus and its relation to depression, anxiety, and stress—a population-based cohort study. Journal of clinical medicine, 12(3), 1169.

Hackett, T., Stepniewska, I., & Kaas, J. (1998). Subdivisions of auditory cortex and ipsilateral cortical connections of the parabelt auditory cortex in macaque monkeys. Journal of Comparative Neurology, 394(4), 475–495.

Hackett, T. A. (2011). Information flow in the auditory cortical network. Hearing research, 271(1-2), 133–146.

Hickok, G., & Poeppel, D. (2007). The cortical organization of speech processing. Nature Reviews Neuroscience, 8(5), 393–402.

Iliadou, V., Bamiou, D.-E., Kaprinis, S., Kandylis, D., & Kaprinis, G. (2009). Auditory Processing Disorders in children suspected of Learning Disabilities—A need for screening? International Journal of Pediatric Otorhinolaryngology, 73(7), 1029–1034.

Jang, S. H., & Choi, E. B. (2022). Evaluation of structural neural connectivity between the primary auditory cortex and cognition-related brain areas using diffusion tensor tractography in 43 normal adults. Medical Science Monitor: International Medical Journal of Experimental and Clinical Research, 28, e936131–936131.

Kaas, J. H. (2008). The evolution of the complex sensory and motor systems of the human brain. Brain research bulletin, 75(2-4), 384–390.

Kanwal, J. S., & Ehret, G. (2010). Communication sounds and their cortical representation. In The auditory cortex (pp. 343-367). Springer.

Karolis, V. R., Corbetta, M., & Thiebaut de Schotten, M. (2019). The architecture of functional lateralisation and its relationship to callosal connectivity in the human brain. Nature communications, 10(1), 1417.

Kim, J.-H., Jung, A.-H., Jeong, D., Choi, I., Kim, K., Shin, S., Kim, S. J., & Lee, S.-H. (2016). Selectivity of neuromodulatory projections from the basal forebrain and locus ceruleus to primary sensory cortices. Journal of Neuroscience, 36(19), 5314–5327.

Köse, B., Karaman-Demirel, A., & Çiprut, A. (2022). Psychoacoustic abilities in pediatric cochlear implant recipients: The relation with short-term memory and working memory capacity. International Journal of Pediatric Otorhinolaryngology, 162, 111307.

Kraus, K. S., & Canlon, B. (2012). Neuronal connectivity and interactions between the auditory and limbic systems. Effects of noise and tinnitus. Hearing research, 288(1-2), 34–46.

Küper, M., Dimitrova, A., Thürling, M., Maderwald, S., Roths, J., Elles, H., Gizewski, E., Ladd, M. E., Diedrichsen, J., & Timmann, D. (2011). Evidence for a motor and a non-motor domain in the human dentate nucleus—an fMRI study. Neuroimage, 54(4), 2612–2622.

Labache, L., Ge, T., Yeo, B. T., & Holmes, A. J. (2023). Language network lateralization is reflected throughout the macroscale functional organization of cortex. Nature communications, 14(1), 3405.

Lee, S. J. (2018). The relationship between hearing impairment and cognitive function in middle-aged and older adults: a meta-analysis. Communication Sciences & Disorders, 23(2), 378–391.

Leong, A. T., Wong, E. C., Wang, X., & Wu, E. X. (2023). Hippocampus Modulates Vocalizations Responses at Early Auditory Centers. Neuroimage, 270, 119943.

Lercher, P., Evans, G. W., & Meis, M. (2003). Ambient noise and cognitive processes among primary schoolchildren. Environment and Behavior, 35(6), 725–735.

Li, X., Yu, K., Zhang, Z., Sun, W., Yang, Z., Feng, J., Chen, X., Liu, C.-H., Wang, H., & Guo, Y. P. (2014). Cholecystokinin from the entorhinal cortex enables neural plasticity in the auditory cortex. Cell research, 24(3), 307–330.

Loughrey, D. G., Kelly, M. E., Kelley, G. A., Brennan, S., & Lawlor, B. A. (2018). Association of age-related hearing loss with cognitive function, cognitive impairment, and dementia: a systematic review and meta-analysis. JAMA otolaryngology–head & neck surgery, 144(2), 115–126.

Lv, X., Wang, Y., Zhang, Y., Ma, S., Liu, J., Ye, K., Wu, Y., Voon, V., & Sun, B. (2024). Auditory entrainment coordinates cortical-BNST-NAc triple time locking to alleviate the depressive disorder. Cell Reports, 43(8).

Malmierca, M. S., & Ryugo, D. K. (2010). Descending connections of auditory cortex to the midbrain and brain stem. In The auditory cortex (pp. 189-208). Springer.

Manno III, F. A., Lau, C., Fernandez-Ruiz, J., Manno, S. H.-C., Cheng, S. H., & Barrios, F. A. (2019). The human amygdala disconnecting from auditory cortex preferentially discriminates musical sound of uncertain emotion by altering hemispheric weighting. Scientific Reports, 9(1), 14787.

Morosan, P., Schleicher, A., Amunts, K., & Zilles, K. (2005). Multimodal architectonic mapping of human superior temporal gyrus. Anatomy and embryology, 210, 401–406.

Munoz-Lopez, M. M., Mohedano-Moriano, A., & Insausti, R. (2010). Anatomical pathways for auditory memory in primates. Frontiers in neuroanatomy, 4, 129.

Nelken, I. (2020). From neurons to behavior: the view from auditory cortex. Current Opinion in Physiology, 18, 37–41.

Nelson, A., & Mooney, R. (2016). The basal forebrain and motor cortex provide convergent yet distinct movement-related inputs to the auditory cortex. Neuron, 90(3), 635–648.

Olavarria, J., & van Sluyters, R. C. (1985). Organization and postnatal development of callosal connections in the visual cortex of the rat. Journal of Comparative Neurology, 239(1), 1–26.

Ong, W., & Garey, L. (1990). Neuronal architecture of the human temporal cortex. Anatomy and embryology, 181, 351–364.

Özkan, M., Köse, B., Algın, O., Oğuz, S., Erden, M. E., & Çavdar, S. (2022). Non-motor connections of the pedunculopontine nucleus of the rat and human brain. Neuroscience Letters, 767, 136308.

Paxinos, G., & Franklin, K. B. (2012). Paxinos and Franklin’s the Mouse Brain in Stereotaxic Coordinates (4th edition ed.). Academic Press.

Perales, M., Winer, J. A., & Prieto, J. J. (2006). Focal projections of cat auditory cortex to the pontine nuclei. Journal of Comparative Neurology, 497(6), 959–980.

Pickles, J. O. (2015). Auditory pathways: anatomy and physiology. Handbook of clinical neurology, 129, 3–25.

Plichta, M. M., Gerdes, A. B., Alpers, G. W., Harnisch, W., Brill, S., Wieser, M. J., & Fallgatter, A. J. (2011). Auditory cortex activation is modulated by emotion: a functional near-infrared spectroscopy (fNIRS) study. Neuroimage, 55(3), 1200–1207.

Puig, M. V., & Gulledge, A. T. (2011). Serotonin and prefrontal cortex function: neurons, networks, and circuits. Molecular neurobiology, 44(3), 449–464.

Remedios, R., Logothetis, N. K., & Kayser, C. (2009). An auditory region in the primate insular cortex responding preferentially to vocal communication sounds. Journal of Neuroscience, 29(4), 1034–1045.

Remedios, R., Logothetis, N. K., & Kayser, C. (2014). A role of the claustrum in auditory scene analysis by reflecting sensory change. Frontiers in systems neuroscience, 8, 44.

Ren, J., Isakova, A., Friedmann, D., Zeng, J., Grutzner, S. M., Pun, A., Zhao, G. Q., Kolluru, S. S., Wang, R., & Lin, R. (2019). Single-cell transcriptomes and whole-brain projections of serotonin neurons in the mouse dorsal and median raphe nuclei. Elife, 8, e49424.

Reznik, D., & Mukamel, R. (2019). Motor output, neural states and auditory perception. Neuroscience & Biobehavioral Reviews, 96, 116–126.

Robert, B., Kimchi, E. Y., Watanabe, Y., Chakoma, T., Jing, M., Li, Y., & Polley, D. B. (2021). A functional topography within the cholinergic basal forebrain for encoding sensory cues and behavioral reinforcement outcomes. Elife, 10, e69514.

Rodgers, K. M., Benison, A. M., Klein, A., & Barth, D. S. (2008). Auditory, somatosensory, and multisensory insular cortex in the rat. Cerebral cortex, 18(12), 2941–2951.

Rönnberg, J., Danielsson, H., Rudner, M., Arlinger, S., Sternäng, O., Wahlin, A., & Nilsson, L.-G. (2011). Hearing loss is negatively related to episodic and semantic long-term memory but not to short-term memory.

Sharma, S., & Bandyopadhyay, S. (2020). Differential rapid plasticity in auditory and visual responses in the primarily multisensory orbitofrontal cortex. Eneuro, 7(3).

Shinn-Cunningham, B. G., & Best, V. (2008). Selective attention in normal and impaired hearing. Trends in amplification, 12(4), 283–299.

Solyga, M., & Keller, G. B. (2025). Multimodal mismatch responses in mouse auditory cortex. Elife, 13, RP95398.

Stansfeld, S. A., Berglund, B., Clark, C., Lopez-Barrio, I., Fischer, P., Öhrström, E., Haines, M. M., Head, J., Hygge, S., & Van Kamp, I. (2005). Aircraft and road traffic noise and children’s cognition and health: a cross-national study. The lancet, 365(9475), 1942–1949.

Stehberg, J., Dang, P. T., & Frostig, R. D. (2014). Unimodal primary sensory cortices are directly connected by long-range horizontal projections in the rat sensory cortex. Frontiers in neuroanatomy, 8, 93.

Teki, S., Kumar, S., von Kriegstein, K., Stewart, L., Lyness, C. R., Moore, B. C., Capleton, B., & Griffiths, T. D. (2012). Navigating the auditory scene: an expert role for the hippocampus. Journal of Neuroscience, 32(35), 12251–12257.

Toga, A. W., & Thompson, P. M. (2003). Mapping brain asymmetry. Nature Reviews Neuroscience, 4(1), 37–48.

Tsukano, H., Hou, X., Horie, M., Kitaura, H., Nishio, N., Hishida, R., Takahashi, K., Kakita, A., Takebayashi, H., Sugiyama, S., & Shibuki, K. (2019). Reciprocal connectivity between secondary auditory cortical field and amygdala in mice. Scientific Reports, 9(1). 10.1038/s41598-019-56092-9

Upadhyay, J., Ducros, M., Knaus, T. A., Lindgren, K. A., Silver, A., Tager-Flusberg, H., & Kim, D.-S. (2007). Function and connectivity in human primary auditory cortex: a combined fMRI and DTI study at 3 Tesla. Cerebral cortex, 17(10), 2420–2432.

Vinogradova, O. (1975). Functional organization of the limbic system in the process of registration of information: facts and hypotheses. In The Hippocampus: Volume 2: Neurophysiology and Behavior (pp. 3-69). Springer.

Vogler, N. W., Chen, R., Virkler, A., Tu, V. Y., Gottfried, J. A., & Geffen, M. N. (2024). Direct Piriform-to-Auditory Cortical Projections Shape Auditory–Olfactory Integration. Journal of Neuroscience, 44(49).

Vogt, B. (2009). Cingulate neurobiology and disease. Oxford University Press, USA.

Wallace, M. T., Ramachandran, R., & Stein, B. E. (2004). A revised view of sensory cortical parcellation. Proceedings of the National Academy of Sciences, 101(7), 2167–2172.

Wang, M., Cao, L., Li, H., Xiao, H., Ma, Y., Liu, S., Zhu, H., Yuan, M., Qiu, C., & Huang, X. (2021). Dysfunction of resting-state functional connectivity of amygdala subregions in drug-naive patients with generalized anxiety disorder. Frontiers in psychiatry, 12, 758978.

Wesson, D. W., & Wilson, D. A. (2010). Smelling sounds: olfactory–auditory sensory convergence in the olfactory tubercle. Journal of Neuroscience, 30(8), 3013–3021.

Winer, J. A. (2010). A profile of auditory forebrain connections and circuits. In The auditory cortex (pp. 41-74). Springer.

Winkowski, D. E., Nagode, D. A., Donaldson, K. J., Yin, P., Shamma, S. A., Fritz, J. B., & Kanold, P. O. (2018). Orbitofrontal cortex neurons respond to sound and activate primary auditory cortex neurons. Cerebral cortex, 28(3), 868–879.

Wise, S., & Jones, E. (1976). The organization and postnatal development of the commissural projection of the rat somatic sensory cortex. Journal of Comparative Neurology, 168(3), 313–343.

Xiao, C., Liu, Y., Xu, J., Gan, X., & Xiao, Z. (2018). Septal and hippocampal neurons contribute to auditory relay and fear conditioning. Frontiers in cellular neuroscience, 12, 102.

Xu, X.-M., Jiao, Y., Tang, T.-Y., Lu, C.-Q., Zhang, J., Salvi, R., & Teng, G.-J. (2019). Altered spatial and temporal brain connectivity in the salience network of sensorineural hearing loss and tinnitus. Frontiers in Neuroscience, 13, 246.

Yao, J. D., Gimoto, J., Constantinople, C. M., & Sanes, D. H. (2020). Parietal cortex is required for the integration of acoustic evidence. Current Biology, 30(17), 3293–3303. e3294.

Yukie, M. (2002). Connections between the amygdala and auditory cortical areas in the macaque monkey. Neuroscience research, 42(3), 219–229.

Zatorre, R. J., Chen, J. L., & Penhune, V. B. (2007). When the brain plays music: auditory–motor interactions in music perception and production. Nature Reviews Neuroscience, 8(7), 547–558.

Zhang, Y., Zhu, M., Sun, Y., Tang, B., Zhang, G., An, P., Cheng, Y., Shan, Y., Merzenich, M. M., & Zhou, X. (2021). Environmental noise degrades hippocampus-related learning and memory. Proceedings of the National Academy of Sciences, 118(1), e2017841117.

Zhong, L., Zhang, Y., Duan, C. A., Deng, J., Pan, J., & Xu, N.-l. (2019). Causal contributions of parietal cortex to perceptual decision-making during stimulus categorization. Nature neuroscience, 22(6), 963–973.

Zhou, X., & Merzenich, M. M. (2012). Environmental noise exposure degrades normal listening processes. Nature communications, 3(1), 843.

